# Quantifying the Morphology and Mechanisms of Cancer Progression in 3D *in-vitro* environments: Integrating Experiments and Multiscale Models

**DOI:** 10.1101/2021.11.16.468856

**Authors:** Nikolaos M. Dimitriou, Salvador Flores-Torres, Joseph Matthew Kinsella, Georgios D. Mitsis

## Abstract

Mathematical models of cancer growth have become increasingly more accurate both in the space and time domains. However, the limited amount of data typically available has resulted in a larger number of qualitative rather than quantitative studies. In the present study, we provide an integrated experimental-computational framework for the quantification of the morphological characteristics and the mechanistic modelling of cancer progression in 3D environments. The proposed framework allows for the calibration of multiscale, spatiotemporal models of cancer growth using state-of-the-art 3D cell culture data, and their validation based on the resulting experimental morphological patterns using spatial point-pattern analysis techniques. We applied this framework to the study of the development of Triple Negative Breast Cancer cells cultured in Matrigel scaffolds, and validated the hypothesis of chemotactic migration using a multiscale, hybrid Keller-Segel model. The results revealed transient, non-random spatial distributions of cancer cells that consist of clustered, and dispersion patterns. The proposed model was able to describe the general characteristics of the experimental observations and suggests that cancer cells exhibited chemotactic migration and accumulation, as well as random motion during the examined time period of development. The developed framework enabled us to pursue two goals; first, the quantitative description of the morphology of cancer growth in 3D cultures using point-pattern analysis, and second, the relation of tumour morphology with underlying biophysical mechanisms that govern cancer growth and migration.

## I. Introduction

### Cancer modelling

Cancer growth and migration are two main characteristics that affect its morphology [1]. Though there is significant knowledge on the qualitative aspects of tumour morphology, its quantitative characterization and the biophysical mechanisms that govern cancer growth and migration remain still elusive. Significant insights into both morphological and mechanistic characteristics of cancer growth can be gained from the use of mathematical models. Spatiotemporal models of cancer growth can be distinguished in three general categories; discrete (e.g. agent based models), continuum (Partial Differential Equations, PDEs), and hybrid models [2]. Each of these categories provides different information on the aspects of tumour growth. Specifically, discrete models can provide information on individual cell processes or tissue microarchitecture [3]. Continuum models have been widely used, initially to describe qualitative aspects of tumour growth, albeit lacking experimental validation [4], and more recently, for a more detailed quantitative description of the macroscopic characteristics of spatiotemporal cancer growth and its response to therapy under both *in-vitro* [5]–[8] and *in-vivo* conditions [9]–[15]. Hybrid models attempt a multiscale description of cancer growth, by incorporating both continuous and discrete variables [16], [17]. Studies from Tweedy et al. [18], [19] utilized experiments and hybrid discrete-continuum (HDC) models of chemotactic migration to investigate the role of self-generated chemotactic gradients in cancer migration. Even though there is a growing literature on spatiotemporal models of cancer, their validation using experimental data is important for quantitatively describing cancer [20]–[28].

### Data acquisition challenges

The validation of a model can be interpreted as the process of quantifying how accurately the predictions describe the experimental measurements [20]. A common reason for the absence of validation in tumour modelling studies is the lack of data availability. *In-vitro* studies usually include the use of 2D cultures [7], [8], resulting in a less realistic representation of cancer growth. *In-vivo* studies, both clinical and experimental, are more realistic; however, they also present limitations. On the one hand, caliper and microCT scan measurements of *in-vivo* tumours in mice do not typically provide information on tumour shape and invasiveness [29], [30]. Intravital imaging is another common way of data collection for *in-vivo* models; however, this technique suffers from technical challenges, such as passive drift of cells or tissues, low penetration depth, tissue heating, and limitations on imaging intervals [1]. On the other hand, clinical data can be limited in terms of time-points [11], resulting in model over-fit.

### 3D cell culture experiments

To this end, 3D cell culture models have become a promising experimental tool. The main reasons are their increased control of the experimental conditions and flexibility of data collection compared to *in-vivo* experiments, as well as their more realistic representation of tumour progression compared to 2D cultures. Differences between 3D and 2D cultures have been observed in cancer growth and its related biochemical processes, such as the secretion of extracellular matrix (ECM) components, and cell-cell interaction components [31], while the histological and molecular features of *in-vitro* 3D spheroids exhibit more similarities to xenografts compared to 2D monolayers [31]. Additionally, the collection of imaging data for *in-vitro* 3D cell cultures is generally easier and more accurate than *in-vivo* models, and high resolution images can be obtained using confocal microscopy. Although 3D cell cultures cannot yet capture the full complexity of tumour growth in a living tissue, overall they yield significant potential for quantitatively describing cancer growth, as they even provide the opportunity to track even single cells.

### Calibration and validation of HDC models

The purpose of the present work is to introduce an integrated framework for the quantitative characterization of spatiotemporal progression of cancer, and its use for multiscale-spatiotemporal model validation for the study of cancer growth mechanisms. The framework presented in Fig. 1 proposes a novel combination of experimental data from state-of-the-art 3D cell cultures, spatial statistical analysis techniques for the quantification of cancer morphology, and a multiscale HDC mathematical model for the quantitative description of the mechanisms underlying cancer progression. Given the spatial scales (μm up to mm) of the 3D cultures, the choice of HDC models instead of purely continuum or discrete models allows us to perform faster calibration on the continuum model component, albeit with a lower fidelity compared to the full model, and validation on the discrete component. In this work, we present a novel approach for model calibration and validation. Instead of splitting the datasets, we perform calibration and validation on the two different levels; calibration on the continuum, and validation on the discrete level. The introduction of the spatial pattern analysis not only enables us to validate the hybrid model, but also to interpret the observed patterns based on the underlying mechanisms.

**Fig. 1:**
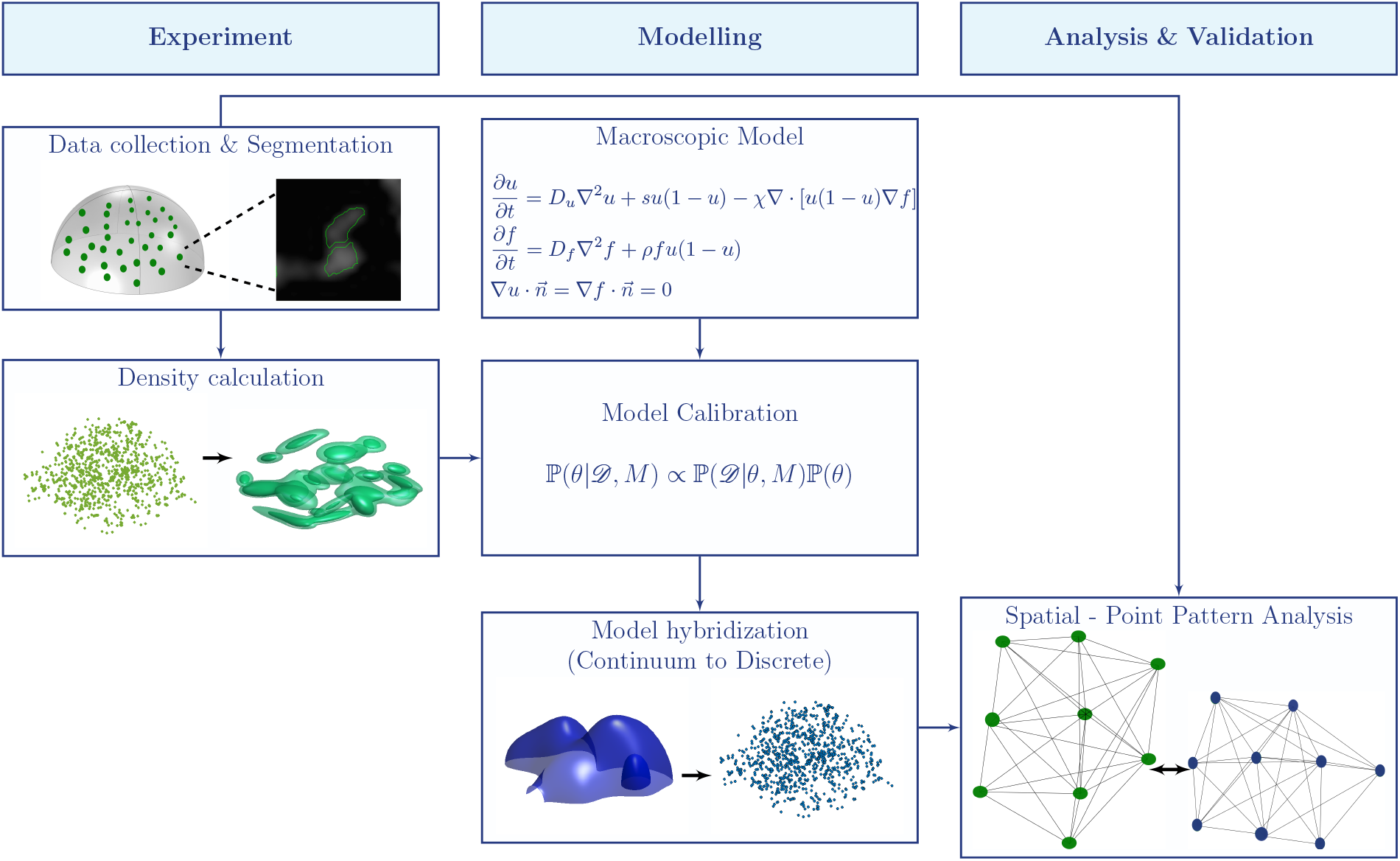
Proposed pipeline for the modelling, validation and analysis of cancer progression using *in-vitro* 3D experimental data.

### The effect of cell sedimentation in 3D cultures

The calibration and validation study of the HDC model was performed using experimental data that present a case of cellular sedimentation, in which the cells were initial distributed across the 3D space and migrated towards the bottom of the space. We attributed this behaviour to a chemotaxis phenomenon based on self-generated gradients, as the 3D environment did not have any imposed chemotactic gradients. Additionally, this phenomenon could not be attributed to gravitational forces, since our calculations showed that the cells were expected to float (Suppl. S.8). This phenomenon has also been observed by Liu et al. [32], and they have interpreted it as the result of cells moving towards the path of least resistance. However, this conjecture disagrees with other observations showing that cells tend to move in the direction of increasing matrix stiffness, termed durotaxis [33].

The rest of the article is organized in a Methods section, where we describe the experiments, data processing, the mathematical model, the calibration and validation techniques, followed by the Results where we present the calibrated model, the validity tests of the full model, as well as a description of the relation between morphology and the underlying mechanisms. Finally, we conclude with the Discussion and Conclusions, where we discuss our results compared to relevant literature, the advantages and limitations of our study, as well as possible extensions and improvements. The code and data of this work are available at https://nmdimitriou.github.io/HyMetaGrowth/.

## II. Methods

### A. Experiments

#### 1) Cell preparation

Triple Negative Breast Cancer (TNBC) cells from the MDA-MB-231 cell line with nuclear GFP (histone transfection), were thawed and cultured at 5% CO2, 37 °C in DMEM (Gibco) at pH 7.2 supplemented with 10% fetal bovine serum (Wisent Bioproducts), 100 U/mL penicillin, 100 μg/mL streptomycin, and 0.25 μg/mL, and amphotericin B (Sigma) in T-75 flasks (Corning). The cells were passaged before reaching 85% confluence. Three passages were performed before the 3D cultures; cells were rinsed twice with DPBS and trypsin-EDTA (0.25%-1X, Gibco) was used to harvest them.

##### 3D cell cultures

A cell-Matrigel (Corning) suspension was created using 0.25 mL of Matrigel (4 °C) and 5 × 10^4^ MDA-MB-231/GFP cells. Droplets of 5 μL cell-Matrigel mixture were deposited onto a high performance glass bottom 6-well plate (0.170 ± 0.005 mm) (Fisher Scientific). The deposited droplets formed a bulk semi-ellipsoid geometry with a uniform distribution of cells across its volume, and separated from each other. In this geometry, we observed vertical migration effects that are attributed to active migration processes as passive migration due to gravitational forces were not sufficient to drag individual cells to the bottom of the space (Suppl. S.8). In total, 12 datasets were produced with 7 samples on days 0, 2, 5, 7, 9, 12, 14 each (Fig. 2). We tracked the viability of the cells by performing flow cytometry (Suppl. S.9).

**Fig. 2:**
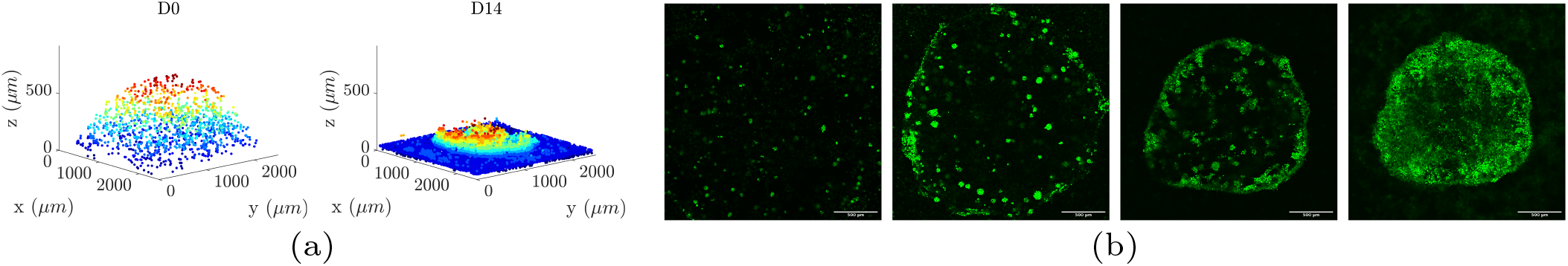
3D cell cultures and segmented nuclei. (a) Segmented cells flowing in 3D on day 0 (left) and sedimented on day 14 (right). (b) Slices at *Z* ≈ 100 μm from the bottom of the plate on days (from left to right) 5, 7, 9, and 14. Scale bar: 500 μm.

#### 2) Imaging and Data preparation

Data acquisition was performed every 2-3 days for a total of 15 days using a confocal microscope (Nikon A1R HD25) coupled with a cell-culture chamber. The dimensions of the 3D cultures were approximately 2.5 × 2.5 × 0.9 mm^3^. Cell localization was made possible by the GFP fluorophore that was present in cell nuclei. The fluorescent nuclei were segmented using an image processing and segmentation pipeline [34]. The preprocessing of the image stacks included: (i) image denoising using the Poisson Unbiased Risk Estimation-Linear Expansion of Thresholds (PURE-LET) technique [35], (ii) intensity attenuation correction across the *z*-dimension [36], (iii) background subtraction using the rolling ball algorithm [37] and manual thresholding of low intensity values using High-Low Look Up Tables (HiLo LUTS), and (iv) cubic spline interpolation of the *xy*-planes of the image stacks. The segmentation of the nuclei was performed using Marker Controlled Watershed segmentation and a classic Distance Based Watershed segmentation to split fused nuclei. The segmented nuclei were then mapped to a 3D Cartesian space by detecting their centroid locations using a 26-connected neighbourhood tracing algorithm implemented in MATLAB [38]. The final step was the calculation of spatial density profiles of the cells represented by their centroids, using the Kernel Density estimation (KDE) via the Diffusion method [39]. KDE was performed using a grid of size (480 × 480 × 176) that matched the spatial grid size of the simulations. The estimated profiles yielded the probabilities of locating the cells in the space. To convert this value into density, we multiplied the probability values at each grid point with the total number of cells, we divided with unit volume, and we corrected by multiplying with the approximate volume of each cell (∼ 3375*µm*^3^).

### B. Multiscale HDC Model

#### 1) Chemotactic hypothesis of cell sedimentation

In our experiment, the cells were initially uniformly distributed in the 3D space, and separated from each other. However, we observed the effect of vertical cell migration leading to sedimentation and aggregation. To model this phenomenon, we hypothesized that this behaviour occurs due to three main reasons; first, the MDA-MB-231 cells are naturally adherent cells, hence the cells tend to remain attached to each other to function properly; second, at the beginning and throughout the course of the experiment, the cells secrete chemotactic signals that enable cell migration and tend to bring the cells closer to each other; third, the cells that are closer to the glass bottom secrete signals at the beginning of the experiment creating a chemotactic gradient decreasing from the bottom towards the top of the space. The rationale behind the third hypothesis is that the glass is a surface that favours cell attachment, hence the cells that are closer to this surface secrete these signals to indicate it as a site of preference. This hypothesis is supported by recent findings on self-generated chemotactic gradients [18], [19], [40], [41] with the difference that we assumed that the chemoattractants stem from the cells and they do not pre-exist in the 3D space.

#### 2) Continuum model

To examine this hypothesis, we used a system of two Keller-Segel (KS) type equations for cancer cell density and chemotactic agent density respectively, which additionally takes into account random motion of cancer cells and chemotactic agents, logistic growth of cancer cells, as well as the increase of chemotactic agents depending on their current concentration in space and the presence of cancer cells. The spatiotemporal evolution of cancer cell, *u*, and chemotactic agent, *f*, densities are obtained by the following PDEs:

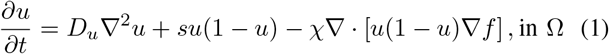

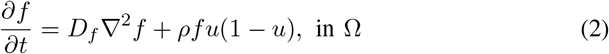

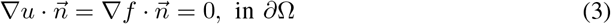

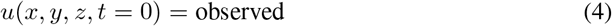

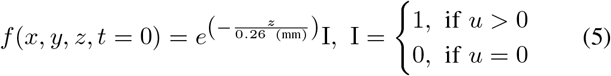

where *D*_*u*_, *D*_*f*_ are the diffusion constants, and *s, ρ* are the growth constants of the cell and signal densities, respectively, and χ is the advection constant of the cells. The right hand side of (1) consists of three terms; the diffusion term *D*_*u*_∇^2^*u* that represents the random motion and expansive growth of the cancer cells, the growth term *su*(1 − *u*) that increases the density of the tumour in a logistic manner, and the nonlinear advection term − χ ∇·[*u*(1− *u*)∇*f*] that represents the biased movement of the cells towards the direction where the gradient of the chemotactic signal density increases. The (1 − *u*) factor in the advection term was added to avoid unwanted overcrowding of the cells that may lead to spikes of cell density [42]. In (2), the evolution of the signal density depends on the diffusion of the signal in 3D space, represented by *D*_*f*_ ∇^2^*f* and the production of signals depending on the current signal density and cell density in space, *ρfu*(1 − *u*). Similarly, (1 − *u*) limits the signal when overcrowding takes place. The spatial domain Ω had the same size as the experimental data, 2.5 × 2.5 × 0.917 mm^3^, and it was represented by 480 × 480 × 176 grid points. We considered no-flux von Neumann boundary conditions (B.C.) in (3), where 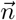 is the outward unit normal to ∂Ω.

The initial conditions (I.C.) for this problem were chosen based on the experimental data and the chemotactic hypothesis. Specifically, the initial cell density profiles of the simulations, (4), were chosen to be the spatial cell density profiles of day 0 of the experiment (Fig. 3). The I.C. for the chemotactic signals were based on the fact that the cells were, initially, uniformly distributed in the 3D space, and separated from each other. Cells attached to the bottom glass were assumed to chemotactic secrete agents first, which in turn promoted the secretion of these agents by the above floating cells as described in (5).

**Fig. 3:**
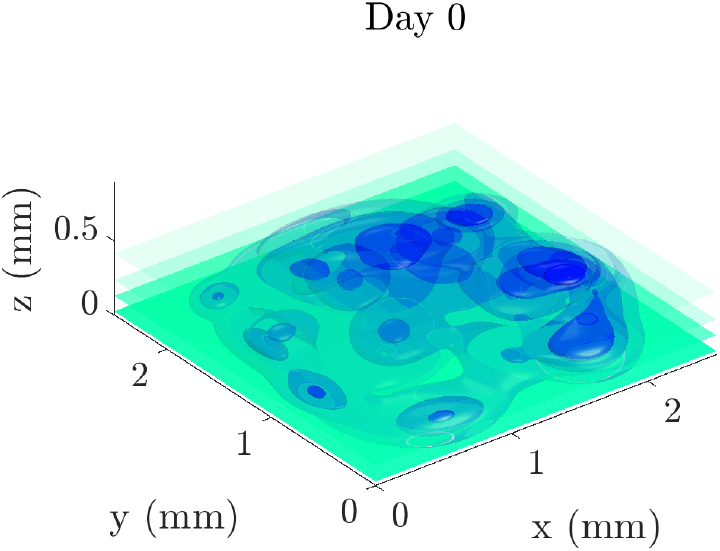
Initial conditions of the continuum model from one of the 12 datasets. The blue colour map represents the cell density profile, *u*, and it is directly obtained from the experimental data. The green colour map represents the chemotactic agents density profile, *f* and it is calculated using (5).

#### 3) Numerical methods

We used the operator-splitting technique to approximate the diffusion, advection, and reaction operators. The diffusion terms were approximated by the Alternating Direction Implicit (ADI) Douglas-Gunn (DG) method [43]. The advection term was approximated by the explicit Lax-Wendroff (LxW) method [44], coupled with the Monotonic Upstream-Centered Scheme for Conservation Laws (MUSCL) flux limiter [45]. The integration in time was performed using the Strang splitting scheme [46]. At every time-step, the Strang splitting scheme evolves the advection and reaction terms by 0.5d*t*, then the diffusion operator by d*t*, and again the advection and reaction operators by 0.5d*t*. The accuracy of this scheme is second-order for both space and time. The proposed numerical scheme was implemented on GPUs using the CUDA/C programming language. Each simulation required approximately 1-5 minutes to complete in a V100-16GB Nvidia GPU (Suppl. S.1).

#### 4) Hybrid model

We hybridized the KS model based on the technique presented in [47], [48]. Specifically, we discretized (1) using the forward time central differences scheme (FTCS) using the approximations found in [49]:

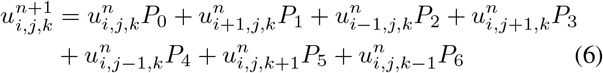

where the grouped terms *P*_*i*_, *i* = 0, …, 6 denote probabilities of the cells of remaining stationary (*P*_0_) or moving back (*P*_1_), front (*P*_2_), left (*P*_3_), right (*P*_4_), down (*P*_5_), up (*P*_6_), defined as

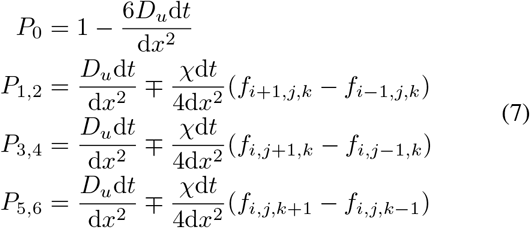

Since the cells were approximately 15 μm in size and the spatial grid points had a distance of 5.2 μm between each other, we assumed that each cell occupied 3 grid points in each direction. To account for this, we modified (6) and (7) by changing the indices that point to a direction to three grid points instead of one, i.e. i±3 instead of i±1 etc. The moving probabilities were then passed to a cellular automaton that updated the position and state of each cell.

The cellular automaton (CA) is presented in Fig. 4a, and its parameters are found in Table I. The CA takes into account three cellular states; alive, quiescent and dead. The CA checks if any cell has reached the proliferation age that is determined based on the estimated parameter *s* (days)^−1^ of the continuum model. We estimated the doubling time from the exponential phase of growth, *e*^*st*^, and the resulting formula *t*_double_ = ln 2*/s*. Considering only the doubling time, would make the cells divide infinitely which is not relevant to the logistic growth dynamics. Hence, we introduced the effect of the space availability, as well as a spontaneous death probability that reduces the viability of the cells. The spontaneous death probability was estimated using experimental data from the later time-points, when the plateau on cell growth takes place. We should note here that some reductions in viability may occur during the exponential phase, however this was already taken into account by estimating *s* from the alive cells only. We tuned the probability based on data obtained from the cell viability assay (Suppl. S.9). If a cell is ready to divide, the algorithm separates into two processes based on cell position in space. If the cell is attached on the glass and there is sufficient space, then the division will be performed on the glass; otherwise, the cell will divide in any direction of the 3D space if there is sufficient space. On the other hand, if there is not sufficient space, the cell becomes quiescent. If the cell is not ready to divide, the CA turns to a migration program.

**Fig. 4:**
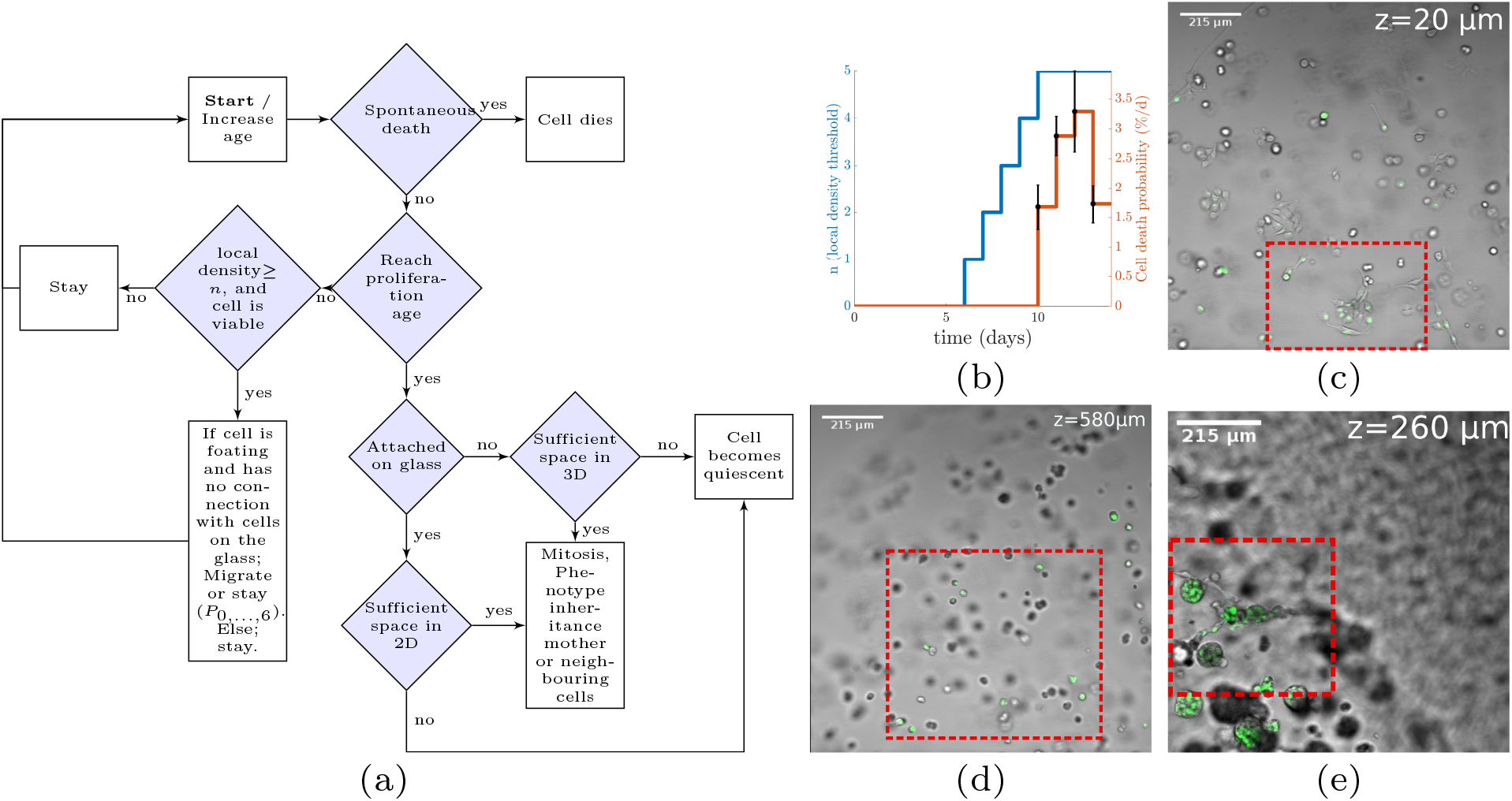
(a) Flowchart of the cellular automaton. (b) Migration, adhesion, and cell death probability parameters of the cellular automaton were changed over time. (c) Cells settled at the bottom (z=20 μm), and (d) cells floating at z=580 μm on day 2. Cells settled on the bottom had stellar shapes, while cells floating in the Matrigel had rounder shapes. (e) Cells floating at z=260 μm on day 7. Some floating cells changed to stellar shapes resembling those attached on the bottom, as shown in panel (c).

**TABLE I:** Parameters and cellular states of the CA.

| Parameters | Units | Interpretation | Value |
| --- | --- | --- | --- |
| $P_{i=0,\dots,6}$ | - | Probability of cell migration and its direction | Calibrated (PDEs) |
| $t_{\text{double}}$ | days | Division time | Calibrated (PDEs) |
| $d_{\text{sp}}$ | days<sup>-1</sup> | Spontaneous death probability rate | Measured |
| $n$ | - | Adhesion parameter for migration | Fixed |
| Cell states |  | Interpretation |  |
| Proliferative |  | Cell has sufficient space to divide. |  |
| Quiescent |  | Cell has not sufficient space to divide. |  |
| Migratory |  | Cell is floating and has sufficient number of neighbours to migrate. |  |
| Non-migratory |  | Cell is attached or connected to the bottom or has not the required number of neighbours to migrate. |  |

The first condition for migration considered an adhesion parameter, and the second is the state of the cell. The adhesion parameter is the local density, defined as the sum of the density in the neighbouring positions, i.e. Σ_*n*=−1,1_(*u*_*i*+*n,j,k*_ + *u*_*i,j*+*n,k*_ + *u*_*i,j,k*+*n*_). A cell can migrate if the local density is equal or greater than a threshold value *n* that is related to the number of neighbouring cells. We hypothesized that the number of neighbours required for cell migration increases over time (Fig. 4b), due to the fact that the initial distribution of cells in the 3D space is sparse; hence they migrate, freely, to search for other cells to attach. However, as cell clustering occurs due to cell division or cell contact, migration becomes less frequent since the cells become more attached to each other. If a cell satisfies these conditions, the algorithm checks the position of the cell. If a cell is settled on the bottom of the space or is connected with a cell located on the bottom, it cannot migrate; otherwise, the cell can migrate in 3D space given the moving probabilities *P*_0_, …, *P*_6_. These two, constrained and unconstrained, migration phenotypes resemble epithelial and mesenchymal phenotypes, respectively, and the transition between them can be found in the literature as mesenchymal to epithelial transition (MET) [50]. Indeed, changes in cellular morphology were observed between cells settled on the bottom and cells floating in the Matrigel. In Fig. 4c, 4d we observe floating cells during the early days of the experiment with round shapes. However, at later time-points (Fig. 4e), we observed stellar shapes for the floating cells probably, due to increased adhesion between them.

### C. Bayesian Inference for calibration of the continuum model

The Keller-Segel model, *M*, (Eq (1)-(3)) includes a set of parameters *θ* = {*D*_*u*_, *s, χ, D*_*f*_, *r*} that are considered unknown. We used their Probability Density Functions (PDF) and the calculated densities from the 3D cell culture data, 𝒟, to assess the most probable parameter values according to Bayes’ rule

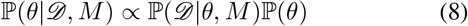

where ℙ (*θ* | 𝒟, *M*) is the posterior PDF of the model parameters *θ* given the observed data 𝒟 and the model *M*, ℙ (𝒟 | *θ, M*) is the likelihood of the observed data 𝒟 given the model *M* and the parameters *θ*, and ℙ (*θ*) is the prior PDF. We assume uninformative, uniform distributions for the model parameter prior PDFs. The experimental data consisted of 12 datasets and each of them had samples collected at 7 time-points. The datasets were assumed to be independent and the model was evaluated for each dataset separately. The likelihood was defined as

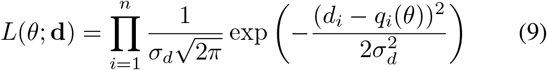

where *n* is the number of spatial grid points, **d** the density profile of the corresponding sample in a dataset, *d*_*i*_, *q*_*i*_ the density values of the experimental sample and simulation result, respectively, at the grid point *i*, and *σ*_*d*_ the variance of the distribution of the likelihood.

We used a Transitional Markov Chain Monte Carlo (TMCMC) algorithm implemented in the Π4U package [51]. The TMCMC algorithm iteratively constructs series of intermediate posterior PDFs

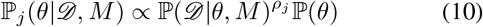

where *j* = 0, …, *m* is the index of the Monte Carlo time series (generation index), and *ρ*_*j*_ controls the transition between the generations, and 0 *< ρ*_0_ *< ρ*_1_ *<* · · · *< ρ*_*m*_ = 1. The TMCMC method can utilize a large number of parallel chains that are evaluated in each Monte Carlo step to reach a result close to the true posterior PDF. To evaluate the calibration results we calculated the Normalized Root Mean Squared Error (NRMSE), the Dice Coefficient (DC), and the cosine similarity, of the density profiles (Suppl. S.4, S.5, S.6).

Since the ratio of model parameters to time-points is small (5:7) for the continuum model, we used all the time-points for the calibration of the continuum model. Validation was performed using the hybrid (discrete-continuum) model using the spatial statistical measures described below.

### D. Spatial Analysis - HDC Model Validation

The introduction of spatial pattern analysis in this work serves two purposes; the quantification of the morphology of cancer, as well as the examination of the validity of the HDC model. The techniques used in this study are described in the following paragraphs.

#### 1) Complete Spatial Randomness Test of Spatial Cell Distributions

The Complete Spatial Randomness (CSR) test examines whether the observed spatial point patterns, in our case the centroids of the nuclei, can be described by a uniform random distribution [52]. The CSR test was performed using Ripley’s *K*-function and the *spatstat* [53] package of R [54]. The *K*-function [55] is defined as the ratio between the number of the events, i.e. locations of points, *j* within a distance *t* from the event *i*, over the total number of events *N*, in the studied volume *V*

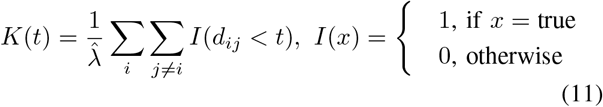

where 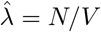 denotes the average density of events, *N*, in the studied volume *V, d*_*ij*_ is the distance between events *i* and *j*, and *t* is the search radius. The *K*-function was calculated for all datasets and compared against complete spatial randomness following a Poisson process *K*(*t*) = 4*πt*^3^*/*3 [55] for three spatial dimensions. Isotropic edge correction was applied in the calculation of the *K*-function. The volume used for the calculation was the same with that used in the simulations, i.e. 2.5 × 2.5 × 0.917 mm^3^. To assess the uncertainty of the random variable *K*, we produced a CSR envelope by generating 100 random distributions and calculating the *K*-function for each of them. The envelope was created by keeping the minimum and maximum values of the resulting *K* values. A substantial upward separation of the observed *K*-function from the theoretical random *K*-function denotes clustered patterns, while a downward separation denotes dispersed patterns [52]. Both separation types suggest non-randomness of the examined spatial distribution.

#### 2) Characterization of the Spatial Cell Distributions

The *Inter-Nucleic (IN) Distance Distribution* for a given sample was calculated by the pairwise Euclidean distances between all nuclei. Given two nuclei *i* and *j* with centroid positions **p**_**i**_ = (*x*_*i*_, *y*_*i*_, *z*_*i*_) and **p**_**j**_ = (*x*_*j*_, *y*_*j*_, *z*_*j*_) respectively, their pairwise Euclidean distance is given by 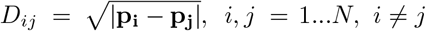, where *N* the total number of nuclei.

The *Nearest-Neighbour (NN) Distance Distribution* for a given sample was calculated using the distances between the nearest neighbours of the nuclei. The nearest neighbour distance for a given nucleus *i* is given by the minimum IN Distance between the nucleus *i* and all the other nuclei of the sample, i.e.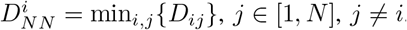.

The comparisons between the *in-vitro* and *in-silico* IN and NN distance distributions were performed using the cosine similarity test [56], in MATLAB [38] (Suppl. S.6).

## III. Results

### A. Sensitivity analysis

To examine the sensitivity of the model parameters, we performed global sensitivity analysis of the model parameters with response to the tumour volume in 3D space, and the tumour area at the bottom with density values greater than 10^−3^. The resulting rank correlation matrix between the model parameters and the outputs confirmed that the signal production rate, *r*, contributed less than the rest of the parameters (Suppl. S.2, Fig. S.4). Thus, we excluded *r* from the calibration process.

### B. Estimation of the macroscopic model parameters

The continuum Keller-Segel model was used to generate simulation data. The resulting cell density profiles for a given parameter set were compared against the *in-vitro* estimated cell density profiles of a dataset. This process was applied to each of the 12 datasets separately. Approximately 14300 different sets of model parameters were assessed using the TMCMC method for each of the 12 datasets. The obtained manifold of the inferred PDFs for one dataset is presented in Suppl. S.3, Fig. S.5a. The marginal distributions and the average values along with their corresponding standard deviations from the posterior PDFs of the model parameters of these datasets are presented in Fig. 5a, and in Suppl. S.3, Fig. S.5b. The estimated model parameters exhibited low uncertainty compared to range of their respective prior PDFs. The growth rate *s* corresponded to a cell doubling time equal to 3.507 ± 0.254 days, (mean ± SEM). The majority of the model parameters are in agreement to each other across different datasets. Some of them, however, varied significantly, suggesting a lack of practical identifiability, i.e. due to small number of time-points. A visual representation of the *in-silico* cell density profiles, presented in Fig. 5b, using the calibrated parameters, shows that the model predictions reproduced the overall behaviour observed in the experiments, i.e. the biased movement of the cells towards the bottom. The Normalized Root Mean Squared Error (NRMSE) of the cell density evaluated at each spatial grid point per time point is presented in Fig. 6, excluding day 0, when the simulation and experimental data were identical. The results show increased errors for days 2 and 14, suggesting that the model was less able to reproduce these two extreme spatial distributions; the highly sparse on day 2, and dense on day 14. The Dice and cosine similarity tests were in agreement to each other, suggesting intermediate similarity between experiments and simulations. The similarity differences can be related to the morphological coherence of the *in-silico* and *in-vitro* datasets, but also to scaffold misalignments among the microscopy sessions, as well as geometric distortions of the scaffold.

**Fig. 5:**
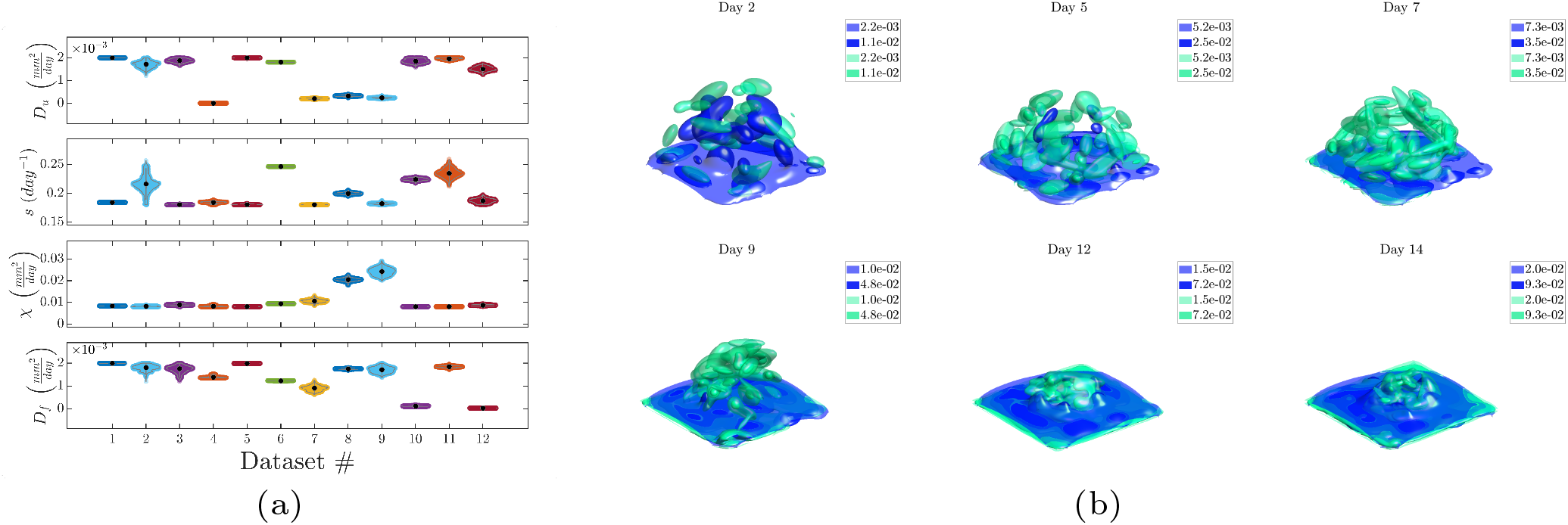
Inferred model parameters and simulation results (a) Violin plots of the marginalized posterior PDFs of the model parameters across the 12 datasets. The black dots represent the median values. (b) Isosurface plot of the experimental and simulated density profiles using the inferred parameters of the initial conditions of a representative dataset. The green colour-map corresponds to the *in-vitro* cell density profiles and the blue colour-map corresponds to the *in-silico* cell density profiles.

**Fig. 6:**
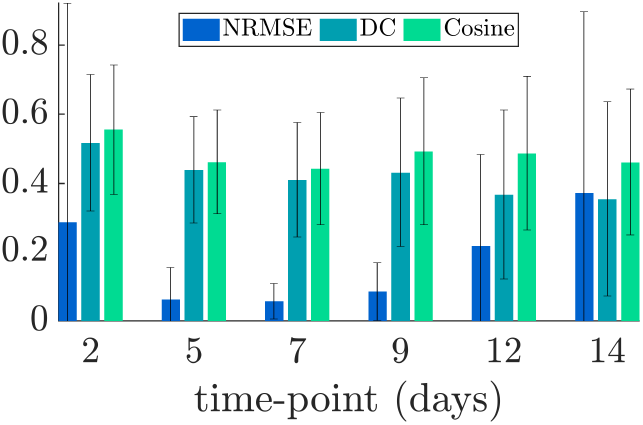
Average and standard deviation of Normalized Root Mean Squared error (NRMSE), Dice coefficient (DC), and cosine similarity across all datasets and across 6 time-points.

### C. Spatial Analysis & HDC Model Validation

The estimated model parameters were subsequently used in the hybrid model (Fig. 4), separately for each dataset. The resulting *in-silico* cellular coordinates were analysed and compared to the corresponding *in-vitro* coordinates of the centroids from the segmented fluorescent nuclei of the cells. The quantitative characterization of the spatial distributions of the cells was performed using the IN, and NN Euclidean distance distributions. The IN distance distributions quantify the positioning of the cells relative to one another, while the NN distance distributions measure the distances between each cell and their nearest neighbouring cell. The resulting IN distance distributions, depicted in Fig. 7a, show that the distributions remained relatively stable across all samples and time, for both experiments and simulations, with a characteristic peak distance at ∼ 1 mm. The cosine similarity test yielded an average similarity value equal to 0.9962 ± 0.0062, suggesting high similarity between IN distance distributions from experiments and simulations. Their similarity remained high across all time-points, as shown in Fig. 7g. On the other hand, the NN distance distributions, presented in Fig. 7b, initially formed wide distributions that gradually tended to become narrower around lower neighbourhood radii values with respect to time, across all samples, with similar characteristic peaks at ∼ 15 μm. These peaks can be interpreted based on the hybrid model hypotheses, specifically regarding the cell division where the daughter cells are placed next to each other, and the adhesion that prevents migration. The average cosine similarity between NN distance distributions from experiments and simulations was equal to 0.7975 ± 0.1457. The similarity between experimental and simulation NN distance distributions decreased as a function of time, as shown in Fig. 7g. We attribute these differences to the variable size of the *in-vitro* cells compare to the fixed size of the *in-silico* cells. According to the definition of NN distance, it can be viewed as a special case of the IN distance. In turn, we would expect that the narrowing of the NN distance distributions would destabilize the IN distance distributions. However, the maintenance of their shape can be interpreted as a result of the organization of the cells into smaller clusters that maintained a relatively constant distance, the synchronized division of the cells, as well as their overall accumulation towards the glass bottom of the wells with respect to time. In other words, the organization of the cells remains similar across space and time.

**Fig. 7:**
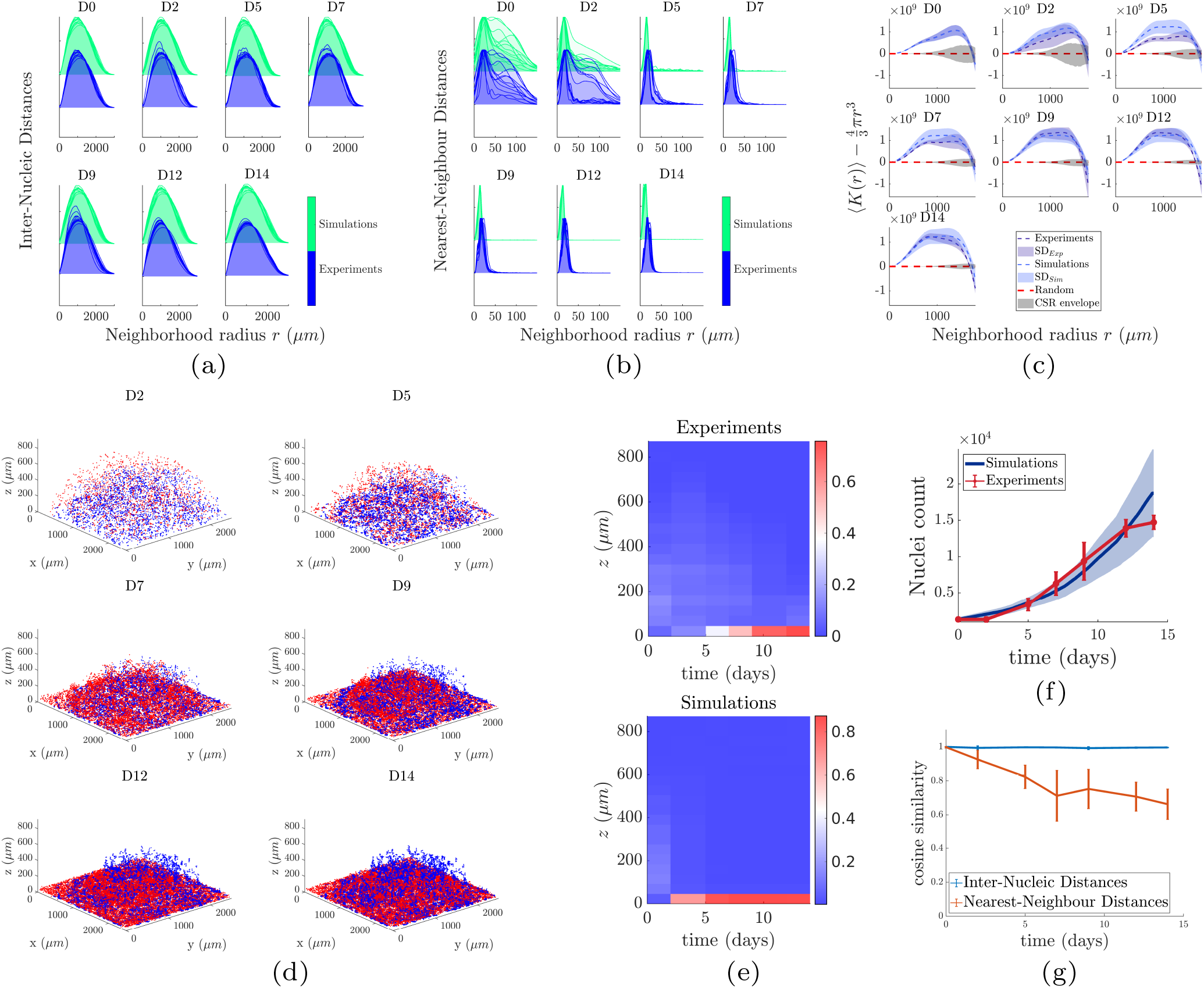
Spatial analysis and comparisons between experiments and simulations (a) Inter-Nucleic Euclidean distance distributions. The title (D#) denotes the time-point in days. (b) Nearest-Neighbour Euclidean distance distributions. (c) Complete Spatial Randomness test; average values of *K*-function across all samples and the corresponding standard error of mean (SEM). (d) Spatial distributions of cells from the cellular automaton (blue) and its corresponding experimental dataset (red) with respect to time. (e) Heatmaps of the normalized number of cells across the *z*-dimension, and across time. The normalization was performed across the z-dimension. (f) Average number of cells across all datasets with respect to time for simulations and experiments. (g) Cosine similarity test for IN and NN distance distributions between experimental and simulation results.

To investigate the spatial organization of the cells, we performed the CSR test, using Ripley’s *K*-function [55]. Specifically, we examined whether the cells, represented by their nuclei centroids, were randomly distributed in space. The results depicted in Fig. 7c indicate substantial differences from a uniform random distribution for both experiments and simulations. For both experimental and simulation data, we observed clustering for a wide range of neighbourhood radii, as well as an increasing dispersion for longer distances across all samples, with respect to time.

The spatial distributions of the *in-silico* cells, and the corresponding experimental dataset are presented in Fig. 7d. We observe that both *in-silico* and *in-vitro* cells performed a biased movement towards the bottom, with the *in-silico* cells characterized also by a more pronounced random motion. Similarly, the numbers of cells across the *z*-dimension in Fig. 7e exhibit similarities between experiment and simulations, even though a relatively small number of cells appears to maintain elevated positions. The *in-vitro* and *in-silico* number of cells with respect to time, shown in Fig. 7f, are in coherence, though the *in-silico* curves did not exhibit a plateau at later stages, suggesting that the spontaneous death probability and the lack of space were not sufficient to stabilize the cell number.

## IV. Discussion

We presented a novel framework that combines 3D cell culture experiments, multiscale models, parameter estimation, and spatial validation techniques to examine and quantify the morphology and mechanisms of cancer progression. We applied the proposed framework to 3D cultures of TNBC cells in Matrigel ECM, and we modeled this behaviour using a multiscale HDC model. The parameters of the continuum model were estimated using Bayesian inference and a TMCMC algorithm. Our results suggest an overall agreement between the calibrated model, and the experimental observations. The NRMSE values suggest that the model performs well on the intermediate time-points, yielding inferior performance for the two extreme cases, i.e. sparse and dense. The DC, and cosine tests suggested that the model exhibits intermediate similarity with the data. However, these differences may have also occurred due to misaligned samples under the microscope or geometric distortions of the scaffolds. The estimated parameters were used in the HDC model for a detailed simulation of the spatial distributions of the cells. The results of both experiments and simulations were analysed using spatial statistical analysis techniques to quantify the morphology of both *in-vitro* and *in-silico* cancer progression. In the following paragraphs, we discuss the relation between the observed spatial patterns, and the underlying mechanisms, particularly as related to the biased movement of the cells towards the bottom.

### A. Relation between Morphological patterns and Biological mechanisms

The continuum KS model consists of diffusion, growth, and advection terms that represent the random motion, proliferation, and biased movement of the cells towards the bottom, respectively. The estimated diffusion constants (Fig. 5a) suggest that random motion together with the unconstrained migration phase affected the distribution of the cancer cells, which was reflected by the increased NN distance values on day 2 (Fig. 7b). The effect of advection, together with the constrained migration were more apparent after day 2 (Fig. 7d). These two parameters reflect the tendency of the cells to form clusters and their tendency to move towards the bottom. This effect was also observed in the NN distances between days 5 and 14 (Fig. 7b), as well as in the heatmaps of the number of cells across different *z*-values (Fig. 7e), which shows a comparable number of *in-silico* and *in-vitro* cells near the bottom. The visualization of the cells (Fig. 7d) shows that the majority of the *in-silico* cells tended to move towards the bottom, while some of them remained floating due to cluster formation. The resulted behaviour lead to a good agreement between the *K*-functions of the experiments and simulations. The cell accumulation resulted in more pronounced clustering patterns for smaller neighbourhood radii with respect to time. The increase in the adhesion parameter with respect to time restricts migration to the cells that have not reached the bottom, contributing to the resulting NN distances. This parameter contributes also to the increased clustering of the *in-silico* cells shown in Fig. 7c, even though the changes in *K*-function were very small.

### B. 3D culture system, biased movement and cell sedimentation

In our experimental setup, we initially mixed cancer cells with Matrigel extracellular matrix (ECM), subsequently deposited droplets of this mixture on a glass-bottom plate, forming semi-ellipsoid scaffolds (Fig. 2), and studied the effects of cell growth for 15 days. The main advantage of this setup is that it allows the observation of the dynamics of the initial phases of cancer growth starting from individual cells, the formation of clusters, and cell migration. In contrast with other 3D spheroid models that require mechanical manipulation (e.g. centrifugation [57]) for the formation of spheroids, possibly causing alterations to intra- and intercellular signalling, the current setup does not require any further manipulation as it allows cells to self-assemble, allowing us to study cell dynamics under more natural conditions. Using this setup, we observed logistic cell growth, cluster formation, as well as, vertical migration towards the bottom starting on Day 2, when most of the cells were still separated from each other. Given the fact that gravity is not sufficient to drag individual cells towards the bottom in a 3D ECM, such as the Matrigel used in the present study (Fig. 2, S.7, Suppl. S.8), we hypothesized that this is a result of active migration regulated by chemosensing, a hypothesis that we examined in this work. Despite the fact that chemotactic migration has been studied on both biological and mathematical levels [58], [59], there is very limited discussion on the observed behaviour of the cells to move towards the bottom of the culture [32].

The selected mathematical model was able to reproduce this biased movement, and the overall framework allowed us to quantify the movement in terms of both spatial patterns and underlying mechanisms. The proposed computational part of the framework allowed us to investigate the mesoscopic scale (μm to mm) taking into account between 1000 and 18000 cells in 3D, exhibiting good performance in terms of processing times. The analysis showed that not all of the *in-silico* cells followed the chemotactic gradient. This phenomenon was also observed by [40], but for a different reason. Their study showed that self-generated gradients may favour the leading wave of cells, because they break down chemoattractants; thus, the cells behind the front do not sense a gradient and move randomly. This phenomenon was not observed in our experiments, due to additional factors that contributed to the biased movement of the cells towards the bottom. These include the compression and degradation of Matrigel, as well as vibrations during the transfer of the samples to the microscope. These factors were not considered in the model; however, the proposed framework provides a promising tool for the study of models of higher complexity.

### C. Splitting calibration and validation on two scales

One of the fundamental limitations of using a HDC model is its increased computational cost. Even though HDC models can provide information on the multiscale mechanisms of cancer growth, their computational cost increases with respect to the number of cells [60]. This problem becomes more apparent during the calibration process that requires many iterations for evaluation of the model parameters. Consequently, some studies [21] fitted the model to the data by performing numerical experiments, which typically is a more qualitative method of model fitting compared to the calibration via maximum likelihood estimation or Bayesian inference. However, this computational cost is usually related to the discrete component of the model. Hence, to overcome this problem we performed calibration only on the continuum part of the model. This process reduced the simulation times to ∼5% of the full model. The calibration process allowed us to estimate the parameters *D*_*u*_, *s, χ* that are then provided to the discrete model. The validation tests were then performed on the full HDC model, even though they were more computationally expensive, they were feasible to implement and provided more details on the predictive performance of the model, as well as a quantified perspective on the morphological behaviour emerging from the examined model.

Typically, the calibration and validation processes are performed in separate datasets. However, since we already had a small number of time-points further data splitting could result in model over-fitting. Another advantage of splitting the calibration and validation tests in two different scales is the preservation of all time-points for both tests. However, the use of all datasets for both calibration and validation may introduce biases. To further examine potential biases introduced in our study, we shuffled the estimated parameters with the datasets, and we calculate the NRMSE for these combinations. The results presented in Suppl. Fig. S.6 show an expected increase in the errors compared to the matched parameter-datasets, the former were roughly found twice as large as the latter, thus indicating the presence of bias, possibly due to variability in the experimentally observed behaviour, as well as the small number of time-points. However, on average the NRMSE remains quite small, while the large standard deviations were present due to increased variance in 2 datasets compared to the rest of them.

### D. Spatial point-patterns as a validity metric for HDC models

The examination of the validity of PDE systems has been well studied; however its application to agent based models (ABMs), including HDC models, has been rather limited. Validation studies on the PDE level typically utilize macroscopic “1-1” relationships between experimental observations and model output, e.g. cell densities, tumour size etc. On the HDC level, this would require measurements that extend to both microscopic and macroscopic levels. As Lima et al. [60] pointed out, obtaining such measurements may be cost-prohibitive or technically infeasible, adding up to the existing problem of sufficient data acquisition. Consequently, some studies evaluated the performance of ABMs using only temporal experimental data [60], [61]. In this work, we approached this challenge on two levels; first, we obtained high quality imaging data such that individual cell segmentation was possible, and second, we utilized this detailed data by performing spatial point-pattern analysis on the discrete cell entities of both the experimental data and the HDC model. The introduction of spatial point-pattern analysis enabled us to utilize the locations of the cells to obtain quantitative information on their organization. The resulting information enabled us to further evaluate the predictive performance of the model taking into account the spatial morphology. In turn, it contributed to the understanding of the spatiotemporal organization of the cells that can further help towards the development of more accurate models.

### E. Conclusion

The proposed framework enabled us to relate the underlying mechanisms of cancer progression with the observed morphological patterns. Future improvements may include incorporating a model term for the quantification of the effect of ECM degradation that may be responsible for the introduction of possible biases. The proposed framework can also be used to study the growth patterns of heterogeneous cell populations such as cancer cells and fibroblasts, as well as, study cancer progression in the presence of therapy. Importantly, potential differences in the morphological patterns in the presence and absence of therapy can be used to design therapeutic strategies that control not only the tumour size, but also their morphological patterns to minimize migration. Overall, the presented framework yields great promise for a more complete quantitative understanding of the organization and progression of cancer.

## ACKNOWLEDGMENTS

N.M.D. thanks the Digital Research Alliance of Canada for the resource allocation RRG # 3975, Ferenc Molnár (University of Notre Dame) for the discussions in GPU computing, Panagiotis Hadjidoukas (IBM Research-Zürich) and Panagiotis Angelikopoulos (D.E. Shaw Research) for the discussions on TMCMC, Remí Dagenais (McGill) for reviewing the article. N.M.D. thanks Stavros Niarchos Foundation (F237055R00), Werner Graupe (F202955R00) and McGill University (90025) for the scholarships. S.F.T. thanks McGill University for the McGill Engineering Doctoral Award (90025) and the FRQNT (291010) for the scholarships.

