## Supplementary for "Quantifying the Morphology and Mechanisms of Cancer Progression in 3D *in-vitro* environments: Integrating Experiments and Multiscale Models"

### Supplementary material

#### S.1 Numerical methods

##### S.1.1 ADI Douglas-Gunn (DG) for the diffusion operator

The ADI-DG scheme [1] is a multi-step method and can be applied to the diffusion term as follows

$$\left(1 - \frac{1}{2}v\delta_x^2\right)u^{n,*} = (1 + \frac{1}{2}v\delta_x^2 + v\delta_y^2 + v\delta_z^2)u^n \tag{1}$$

$$\left(1 - \frac{1}{2}v\delta_y^2\right)u^{n,**} = u^{n,*} - \frac{1}{2}v\delta_y^2 u^n \tag{2}$$

$$\left(1 - \frac{1}{2}v\delta_z^2\right)u^{n+1} = u^{n,**} - \frac{1}{2}v\delta_z^2 u^n \tag{3}$$

where, $v = \frac{D_u \mathrm{dt}}{2h^2}$, $h$ is the spatial grid step, assuming $h = \mathrm{dx} = \mathrm{dy} = \mathrm{dz}$, $\delta^2_{x,\ y,\ z}$ are the central difference operators for the second derivatives in $x,\ y,\ z$ respectively, and $u^{n,*},\ u^{n,**}$ are the intermediate values of $u$.

To examine the performance of the ADI-DG scheme we performed some test simulations on the simple 3D diffusion model with von Neumann boundary conditions,

$$\frac{\partial u}{\partial t} = D\nabla^2 u \text{ in } \Omega, \tag{4}$$

$$\frac{\partial u}{\partial \vec{n}} = 0 \text{ in } \partial\Omega \tag{5}$$

and we compared these results with the explicit 7-point and 19-point [2] central differences. The results for a line across the 3D space are presented in Fig. S.1, showing similar behaviour across the 3 numerical methods. The advantage of using the ADI-DG scheme is that it is unconditionally stable, hence we can use a larger temporal grid size compared to the explicit central differences. However, the step size should be selected with caution since a large step size may lead to inaccuracies in the transient behaviour. In section S.1.3 we provide the step size used for this study.

##### S.1.2 Explicit Lax-Wendrof (LxW) with Monotonic Upstream-Centered Scheme for Conservation Laws (MUSCL) flux limiter for the advection operator

The explicit LxW-MUSCL [3, 4] method applied to the advection term can be written as follows

$$u^{n+1}_{i,j,k} = u^n_{i,j,k} + \chi\frac{\mathrm{d}t}{h}(F_{i-1/2} - F_{i+1/2} + F_{j-1/2} - F_{j+1/2} + F_{k-1/2} - F_{k+1/2}) \tag{6}$$

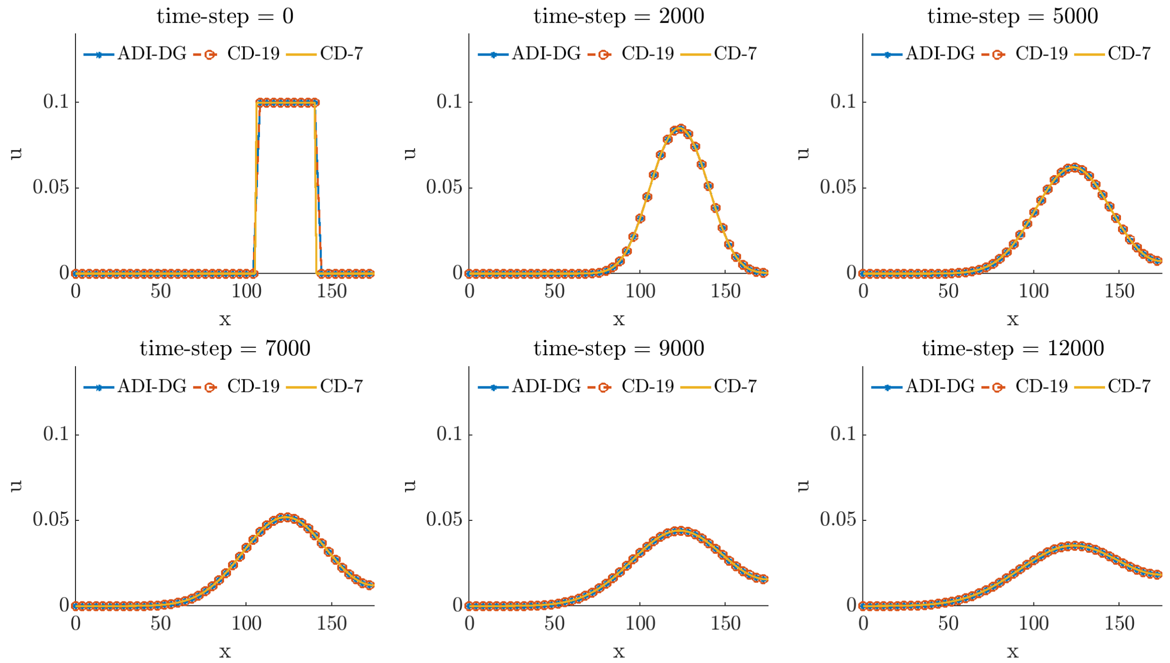


Figure S.1: Performance of the ADI-DG scheme compared to the 7-point and 19-point central differences. The results correspond to density profiles on a line of the 3D space. As expected the 3 methods behave in a similar manner.

Here, $F_{i\pm1/2}$ are defined as follows

$$F_{i-1/2} = (u\nabla f)_{i-1} + \phi_{-}\frac{1}{2}\mathrm{sign}((\nabla f)_i)(1-c)[u_i(\nabla f)_i - u_{i-1}(\nabla f)_{i-1}] \tag{7}$$

$$F_{i+1/2} = (u\nabla f)_{i} + \phi_{+}\frac{1}{2}\mathrm{sign}((\nabla f)_i)(1-c)[u_{i+1}(\nabla f)_{i+1} - u_{i}(\nabla f)_{i}] \tag{8}$$

where $c = \chi\frac{\mathrm{dt}}{h}$,

$$(u\nabla f)_i = u_i \max(0, (\nabla f)_i) - u_{i+1}\max(0, -(\nabla f)_{i+1}) \tag{9}$$

$$(u\nabla f)_{i-1} = u_{i-1} \max(0, (\nabla f)_{i-1}) - u_{i}\max(0, -(\nabla f)_{i}) \tag{10}$$

and

$$(\nabla f)_i = \frac{f_i - f_{i-1}}{h}, \quad (\nabla f)_{i-1} = \frac{f_{i-1} - f_{i-2}}{h}, \quad (\nabla f)_{i+1} = \frac{f_{i+1} - f_{i}}{h} \tag{11}$$

The $\phi_{\pm}$ are the flux limiter variables and are defined as follows

$$\phi_{\pm} = \phi(r_{i\pm1/2}) = \max(0, \min(2r_{i\pm1/2}, \frac{1}{2}(r_{i\pm1/2}+1), 2)) \tag{12}$$

where,

$$r_{i-1/2} = \frac{u_I - u_{I-1}}{u_i - u_{i-1}}, \; r_{i+1/2} = \frac{u_{I+1} - u_I}{u_{i+1} - u_i} \tag{13}$$

and $I = i - \mathrm{sign}((\nabla f)_i)$. The same procedure is repeated for $F_{j\pm1/2}$ and $F_{k\pm1/2}$. The LxW-MUSCL method is conditionally stable. Hence, the size of the time-step should satisfy the Courant-Friedrichs-Lewy (CFL) condition, $\mathrm{d}t \leq \frac{1}{3}\frac{h}{\chi \max((\nabla f)_{i,j,k})}$

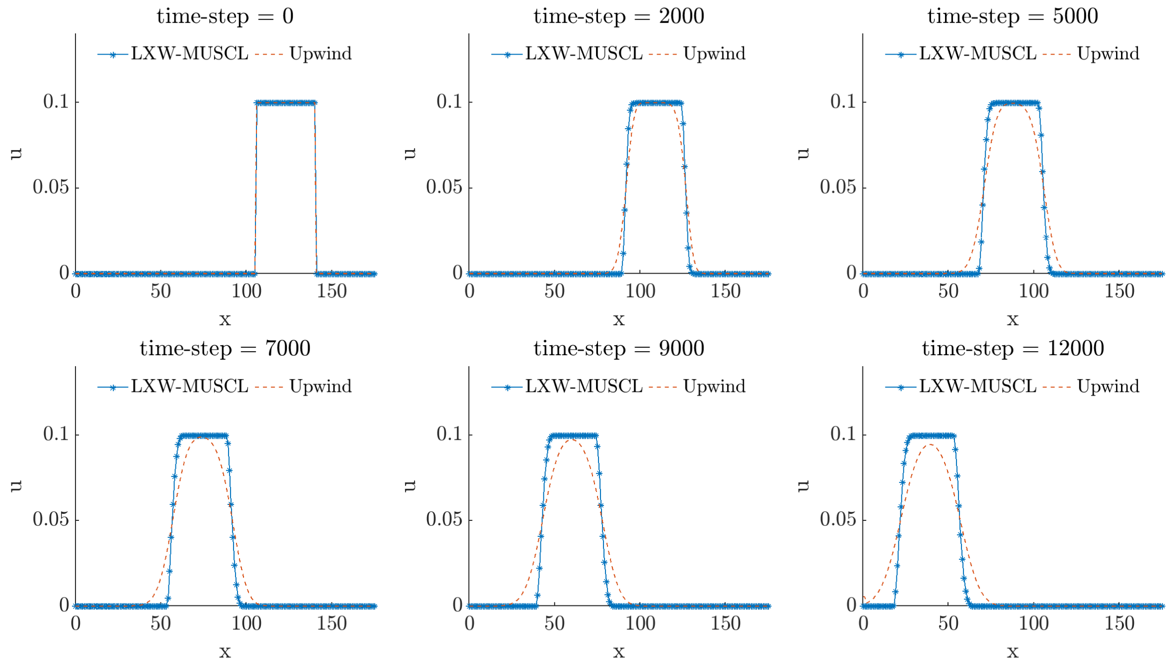


Figure S.2: Performance of the LxW-MUSCL scheme compared to the 1st order upwind scheme. Both methods are applied to the advection equation of (14). The LxW-MUSCL scheme is able to preserve the shape of the waveform as expected from the advection equation. The upwind scheme exhibits a diffusive behaviour as time increases, a result that does not follow the advection equation.

To examine the performance of the LxW-MUSCL scheme we performed some test simulations on the simple 3D advection model with von Neumann boundary conditions,

$$\frac{\partial u}{\partial t} = -c\nabla \cdot u \text{ in } \Omega, \tag{14}$$

$$\frac{\partial u}{\partial \vec{n}} = 0 \text{ in } \partial\Omega \tag{15}$$

and we compared these results with the 1st order upwind scheme. The results for a line across the 3D space are presented in Fig. S.2, and show that the LxW-MUSCL scheme is able to preserve the wave form across time. In contrast, the upwind scheme diffuses in time, a result that is not expected from the advection term, indicating the advantage of using the LxW-MUSCL scheme.

#### S.1.3 Strang splitting

The Strang splitting technique [5] to preserve second-order accuracy in time is the following; For each time-step:

1. evolve the explicit term (advection) for a time-step $\mathrm{d}t/2$,
2. evolve the implicit term (diffusion) for a time-step $\mathrm{d}t$,
3. evolve the explicit term (advection) for a time-step $\mathrm{d}t/2$.

Since we use two numerical methods; an implicit and an explicit, we can use two different time steps. The LxW-MUSCL method is conditionally stable and we choose a time-step based on the CFL condition $\mathrm{d}t \leq \frac{h}{\chi \max((\nabla f)_{i,j,k})}$. The time-step is adaptive and has to be re-evaluated in every time-step, because it depends on $\max((\nabla f)_{i,j,k})$, which is also also dynamic. On the other hand, the ADI-DG is unconditionally stable, hence a larger time-step can provide sufficiently

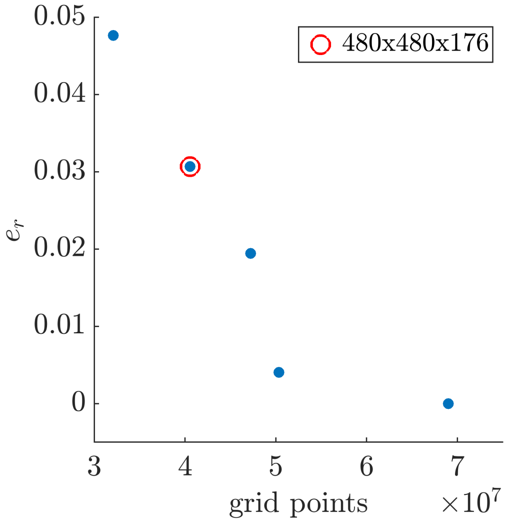


Figure S.3: Grid size error analysis. The relative error of the density integral converges as the grid size increases. The selected grid size has approximately 3% difference from the finest grid.

smooth solutions. We selected the time-step for the ADI-DG as $h^2/\max(D_u, D_f)$, a value that would violate the von Neumann stability condition in the case of an explicit central differences scheme, but not too large to yield inaccuracies in the transient behaviour. The two different time-steps require a modification of the Strang splitting to the following form; For each implicit time-step:

1. evolve the explicit term for j steps until $\sum_j \mathrm{d}t_{\mathrm{exp}} = \mathrm{d}t_{\mathrm{imp}}/2$,
2. evolve the implicit term for $\mathrm{d}t_{\mathrm{imp}}$,
3. evolve the explicit term for j steps until $\sum_j \mathrm{d}t_{\mathrm{exp}} = \mathrm{d}t_{\mathrm{imp}}/2$.

where $\mathrm{d}t_{\mathrm{exp}}$ and $\mathrm{d}t_{\mathrm{imp}}$ are the time-steps for the LxW-MUSCL and ADI-DG methods, respectively.

The implementation of the numerical schemes was performed in CUDA/C language [6]. More information on implementation can be found on the documentation of the code available at `https://nmdimitriou.github.io/HyMetaGrowth/`.

#### S.1.4 Grid size error analysis

To examine the numerical errors introduced by the size of the spatial grid, we performed several simulations with variable grid size and the same initial conditions. The results for five different grid sizes, including the grid size used for this study, show that the errors converge as the grid size increases (Fig. S.3). The relative error between the selected grid and the finest examined grid is approximately 3%. The relative error, $e_r$, was calculated on the integral of the density in the 3D space for every grid, $c_g$, and having as reference the results from the finest grid, $c_f$, as shown in (16), where $u$, (17), is a matrix containing the cell density at each grid point.

$$e_r = \frac{|c_g - c_f|}{c_f}, \tag{16}$$

$$c_i = \int u \mathrm{d}V \tag{17}$$

### S.2 Global sensitivity analysis

The results obtained from the calibration of the continuum spatiotemporal model against the experimental data using TMCMC algorithm indicated increased variability of the chemotactic signal production rate, $r$, across the samples. This

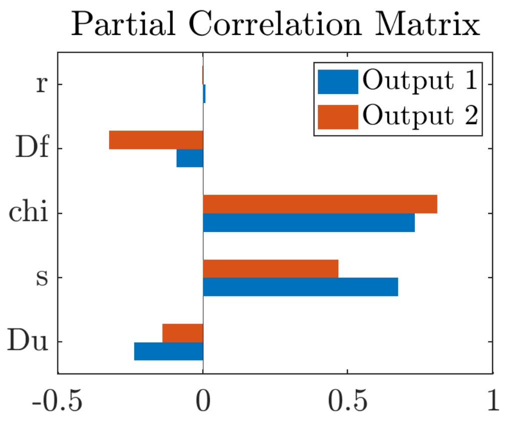


Figure S.4: Global sensitivity analysis of the model parameters with respect to the tumour volume with density values greater than $10^{-3}$ (Output 1), and area of tumour found at the bottom $xy-$plane with density values greater than $10^{-3}$ (Output 2). The partial correlation matrix shows that the chemotactic signal production rate, $r$, was less correlated to the output. Hence it exhibited less contribution compared to the rest of the parameters.

may indicate decreased sensitivity of the parameter $r$ to the final output. To examine this hypothesis, we performed global sensitivity analysis for the model parameters with respect to the model output. The sensitivity analysis was performed using the Latin Hypercube Sampling (LHS) method [7]. The model parameters (input) were examined with respect to the tumour volume with density values greater than $10^{-3}$ (Output 1), as well as the area of tumour found at the bottom of the space with density values greater than $10^{-3}$ (Output 2). The sampled parameter space remained the same as the one used for the calibration. In total, 512 samples were produced, and the relation of the model response to the input was calculated using the Partial Rank Correlation Matrix between input and outputs 1 and 2. The results depicted in Fig. S.4 denote that the chemotactic signal production rate, $r$ was less correlated to both outputs, making $r$ less sensitive compared to the rest of the parameters. Additionally, we observed that the advective constant, $\chi$, and the tumour growth rate, $s$, were positively correlated to both outputs, hence they contributed to tumour clustering. In contrast, the diffusion constants $D_f$, $D_u$ were negatively correlated with both outputs, and they contributed to the dispersion of the tumour. Both observations are consistent with respect to the function of the model.

### S.3 Continuum model calibration

The obtained manifold of the inferred PDFs of the model parameters for one dataset using the TMCMC method is presented in Fig. S.5a. In Fig. S.5b we observe the changes of the chemotactic signalling gradient at two different. The gradient became less steep with respect to time, thus making chemotactic migration less pronounced at later time-points, compared to earlier time-points.

### S.4 Normalized Root Mean Squared Error

The differences between the *in-vitro* and *in-silico* estimated cell density profiles were calculated using the Normalized Root Mean Squared Error (NRMSE) that is defined in (18), where $u_e$, $u_s$ are the arrays containing the experimental and simulated density profiles, respectively.

$$\text{NRMSE} = \frac{1}{\max(u_e) - \min(u_e)} \sqrt{\frac{\sum_{i=1}^{N}(u_s^i - u_e^i)^2}{N}} \tag{18}$$

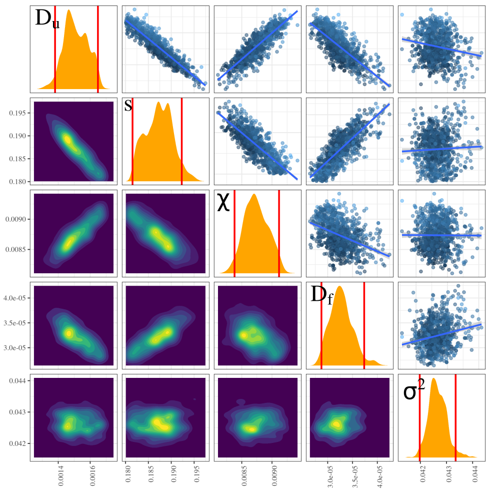


(a)

| Dataset | $D_u \in [10^{-3}, 2] \times 10^{-3}$ $(mm^2 d^{-1})$ | $s \in [1.8, 3] \times 10^{-1} (d^{-1})$ | $\chi \in [0.8, 5] \times 10^{-2} (mm^2 d^{-1})$ | $D_f \in [10^{-3}, 2] \times 10^{-3}$ $(mm^2 d^{-1})$ |
|---|---|---|---|---|
| 1 | $1.999 \pm 0.001$ | $1.843 \pm 0.001$ | $0.833 \pm 0.001$ | $1.992 \pm 0.003$ |
| 2 | $1.698 \pm 0.138$ | $2.164 \pm 0.140$ | $0.819 \pm 0.014$ | $1.792 \pm 0.120$ |
| 3 | $1.872 \pm 0.070$ | $1.805 \pm 0.003$ | $0.883 \pm 0.028$ | $1.724 \pm 0.149$ |
| 4 | $0.006 \pm 0.001$ | $1.841 \pm 0.017$ | $0.830 \pm 0.028$ | $1.404 \pm 0.050$ |
| 5 | $1.995 \pm 0.004$ | $1.806 \pm 0.005$ | $0.802 \pm 0.001$ | $1.983 \pm 0.006$ |
| 6 | $1.807 \pm 0.001$ | $2.463 \pm 0.001$ | $0.936 \pm 0.001$ | $1.229 \pm 0.001$ |
| 7 | $0.205 \pm 0.014$ | $1.803 \pm 0.002$ | $1.063 \pm 0.068$ | $0.895 \pm 0.108$ |
| 8 | $0.328 \pm 0.026$ | $1.998 \pm 0.016$ | $2.050 \pm 0.069$ | $1.748 \pm 0.019$ |
| 9 | $0.244 \pm 0.022$ | $1.823 \pm 0.011$ | $2.414 \pm 0.166$ | $1.703 \pm 0.124$ |
| 10 | $1.854 \pm 0.085$ | $2.243 \pm 0.020$ | $0.803 \pm 0.002$ | $0.118 \pm 0.019$ |
| 11 | $1.951 \pm 0.035$ | $2.350 \pm 0.073$ | $0.804 \pm 0.003$ | $1.840 \pm 0.034$ |
| 12 | $1.514 \pm 0.090$ | $1.871 \pm 0.036$ | $0.874 \pm 0.024$ | $0.032 \pm 0.003$ |

(b)

Figure S.5: Model Calibration (a) Model parameter inference for 1 of the 12 datasets using the TMCMC method. Above diagonal: Projected TMCMC samples of the posterior distribution in 2D space. Diagonal: Marginals of the joint posterior obtained via kernel densities. Red lines denote the 95% credible intervals. Below diagonal: 2D projected densities of the posterior obtained using 2D kernel densities. (b) Average and standard deviation of the inferred parameter values of the continuum KS model across all datasets. The prior PDF boundaries used for the parameter estimation are shown in the header of the table.

### S.5 Dice coefficient

In addition to the NRMSE, we used the Dice coefficient to compare the *in-vitro* and *in-silico* cell density profiles. Given two vectors, $\vec{A}$, $\vec{B}$ the Dice coefficient is defined as

$$\mathrm{DC} := \frac{2\vec{A} \cdot \vec{B}}{||\vec{A}||_2 + ||\vec{B}||_2} \tag{19}$$

where $|| \cdot ||_2$ the Euclidean norm. A Dice coefficient of 1 denotes total overlap between the two vectors.

### S.6 Cosine similarity test

The *in-vitro* and *in-silico* cell density profiles, as well as the IN Distance Distributions, were compared using the cosine similarity measure [8]. The cosine similarity measure emerges from the Euclidean dot product, whereby the similarity of two given vectors $\vec{a}$, $\vec{b}$, is defined as

$$\mathrm{sim}(\vec{a}, \vec{b}) := \cos(\vec{a}, \vec{b}) = \frac{\vec{a} \cdot \vec{b}}{||\vec{a}||_2 ||\vec{b}||_2} \tag{20}$$

where $|| \cdot ||_2$ the Euclidean norm. The similarity measure can take values between -1 and 1 indicating exactly opposite and identical vectors respectively. A similarity value of zero denotes orthogonal vectors. Increasing values between 0 and 1 denote low, intermediate and high similarity.

### S.7 Biases introduced by the calibration and validation

The preservation of all time-points for both calibration and validation may have introduced some biases in the model parameter estimates. To quantify these biases of the proposed method, we calculated the NRMSEs using different combinations of parameter sets and experimental datasets. The total number of evaluations is given by

$$12 \text{ parameter sets} \times 12 \text{ datasets} - 12 \text{ matched parameters-datasets} = 132$$

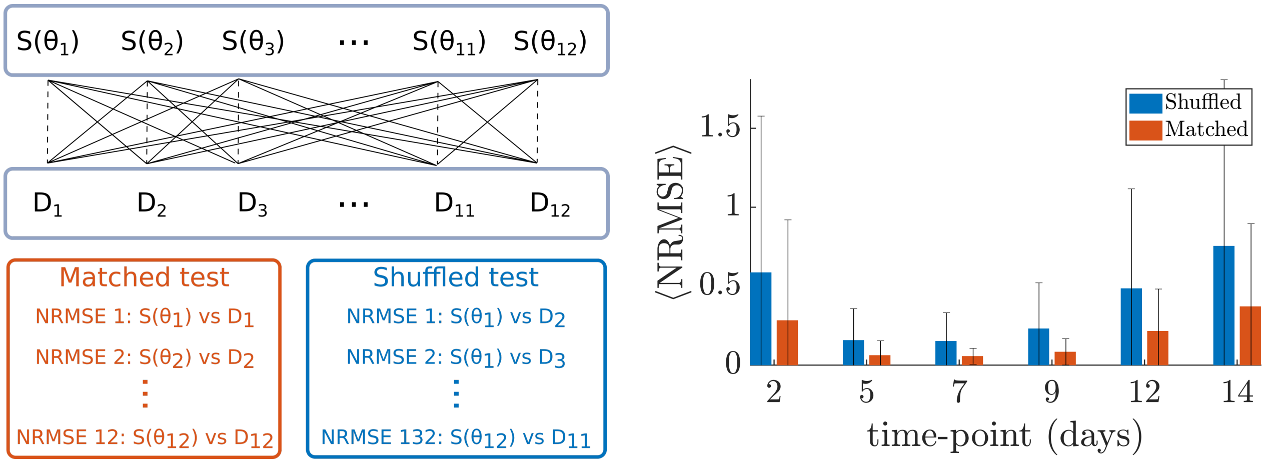


(a) (b)

Figure S.6: Estimation of bias for the calibration and validation framework. (a) Schematic representation of the shuffling technique. Simulations using parameter sets estimated from their corresponding dataset were compared against a different dataset. Here, $\theta_i$ are the parameter sets estimated from the experimental dataset $D_i$, and $S(\theta_i)$ is the simulation output resulting from the parameter set $\theta_i$. The bias estimation was performed using the combinations of simulations and datasets that are connected with continuous lines. The parameter sets estimated from their corresponding datasets are connected with dashed arrows and were not taken into account in the bias estimation. (b) The errors obtained from the shuffled tests were higher compared to those in the matched test, but remain within acceptable levels.

evaluations. The results presented in Fig. S.6 show an increase in the shuffled NRMSEs, which were found to be approximately twice as high as in the matched parameters-datasets. The errors for half of the examined time-points remained at relatively low levels.

### S.8 The effect of gravity on the 3D culture

We examined the effect of gravitational forces in the biased movement and accumulation of the cells at the bottom of the 3D cultures. Specifically, we calculated the gravitational force on a cell under the assumption of spherical shape, and a weight adopted from [9]. We compared the magnitude of the gravitational force with that of the force exerted by Matrigel in a cell as shown in Fig. S.7. The resulting forces were found to be comparable, indicating the cell aggregation at the bottom is not likely a result of gravity.

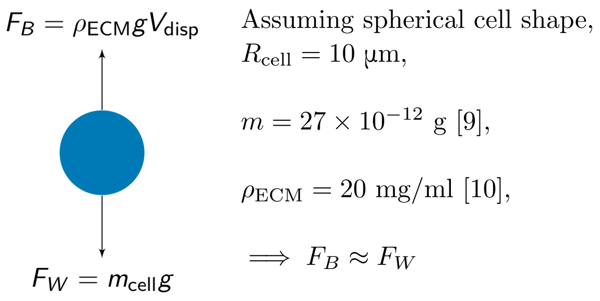


Figure S.7: Gravity in the 3D culture (a) Schematic representation of a cell and the two opposing forces exerted on it (b) The gravitational and its opposite force are approximately equal, thus suggesting that gravity was likely not responsible for the observed sedimentation.

### S.9 Cell viability assay

To estimate the viability of the cells during the plateau phase of the logistic curve, we performed a cell viability test on the 3D cultures using flow cytometry. The number of dead cells was measured for days 11, 12, and 13 after the time of seeding. For each sample, the following protocol was implemented. Time 0 was acquired from non-encapsulated cells and every other time point corresponds to cells extracted from Matrigel domes. Fresh cell culture media was warmed at 37°C and supplemented with collagenase and dispase 1X (Sigma Aldrich, Cat. # 11097113001). Micropipettes were used to break down the cell laden Matrigel constructs by suction force, and these were incubated for 30 minutes inside a cell culture incubator (95% relative humidity, 37°C, and 5% $CO_2$). Cells were centrifuged at 500g for 5 minutes, the supernatant was discarded and a 0.05% trypsin (Wisent, Cat. # 325-042-CL) was used to break any cell-to-cell adhesions for a period of 5 min. Trypsin was neutralized with cell culture medium, and the cell pellet was recollected after 5 min of centrifugation at 500g. A solution containing warm (37°C) 1X DPBS and 1X propidium iodine (miltenyi biotech, Cat. # 130-093-233) was used to resuspend and incubate the collected cells for 15 minutes. Finally, cells were centrifuged again at 500g for 5 minutes, the supernatant was discarded and they were resuspended in DPBS prior to flow cytometry experimentation. All time-points were acquired in triplicates and a PI+ control was implemented using fixed (4% PFA, 5 min) and pierced cells (0.01% SDS, 1 min). Dead cells were identified using the gating strategy described in Crowley et al. [11]. The percentage of dead cells was used to directly obtain the cell death probability rate, which is presented in Fig. 4b of the manuscript.
